# CoCoNet: Boosting RNA contact prediction by convolutional neural networks

**DOI:** 10.1101/2020.07.30.229484

**Authors:** Mehari B. Zerihun, Fabrizio Pucci, Alexander Schug

**Author notes:** Contributed equally to this work (Mehari B. Zerihun, Fabrizio Pucci).

## Abstract

Physics-based co-evolutionary models such as direct coupling analysis (DCA) in combination with machine learning (ML) techniques based on deep neural networks are able to predict protein contact maps with astonishing accuracy. Such contacts can be used as constraints in structure prediction and massively increase prediction accuracy. Unfortunately, the same ML methods cannot readily be applied to RNA as they rely on large structural datasets only available for proteins but not for RNAs. Here, we demonstrate how the small amount of data available for RNA can be used to significantly improve prediction of RNA contact maps. We introduce an algorithm called **CoCoNet** that is based on a combination of a **Co**evolutionary model and a shallow **Co**nvolutional Neural **Net**work. Despite its simplicity and the small number of trained parameters, the method boosts the contact prediction accuracy by about 70% with respect to straightforward DCA as tested by cross-validation on a dataset of about sixty RNA structures. Both our extensive robustness tests and the limited number of parameters allow the generalization properties of our model. Finally, applications to other RNAs highlight the power of our approach. CoCoNet is freely available and can be found at https://github.com/KIT-MBS/coconet.

## 1 Introduction

Ribonucleic acid (RNA) is one of biomolecular key players in cells by playing significant roles in many biological activities such as the coding, regulation and expressions of genes. For examples, non-coding RNA is involved in genetic regulation acting on transcriptional and translational machineries [1, 2] thus enables life as we know it. Since RNA function is closely related to its three-dimensional (3D) structure, experimental techniques such as X-ray diffraction and nuclear magnetic resonance (NMR) are the methods of choice to experimentally determine RNA 3D structure. However, these approaches can be very challenging for RNA that is characterized by a high conformational flexibility. This is reflected in the limited number of RNA 3D structures in the Protein Data Bank (PDB) representing only few percents of the total number of all PDB entries [3]. The large majority of known RNAs remain thus still structurally unresolved and is sometimes even called the dark matter of the biomolecular universe [4].

Computational methods can be a powerful tools to complement experimental efforts by predicting and analyzing RNA structures and can be used alone or in combination with experimental and statistical methods. When direct structure determination is not feasible, indirect measurement might still be possible. To improve the interpretation of such indirect experimental data, they can be integrated in computational modeling tools. For instance, small angle X-ray scattering (SAXS), and single molecule Förster Resonance Energy Transfer (FRET) data have been fruitfully used in combination with molecular dynamics simulations of proteins [5, 6]. Similarly, homology modeling, fragment- and physics-based structure prediction approaches have been developed in the last decade [4, 7, 8, 9, 10, 11, 12, 13, 14, 15] and their accuracy and efficiency, while remain limited especially for large RNAs, is constantly improving as shown in the four blind prediction experiments RNAPuzzle [16, 17, 18, 19].

Likewise, information about spatial proximity of nucleotides inferred by statistical approaches from multiple sequence alignment (MSA) of RNA families can be utilized as spatial constrains in molecular modeling tools [4, 20, 21, 22]. Since structure prediction methods in tandem with these prior information have shown to be more accurate than used alone, these statistical methods have received lot of attention. A wide range of methods based on different implementations of direct coupling analysis (DCA) [23, 24] of coevolving nucleotides, including the mean-field approximation, pseudo-likelihood maximization, sparse inverse covariance estimation and Boltzmann learning [25, 26, 27, 28, 29, 30, 31], have been thus recently introduced to improve the reliability of predicting nucleotides sharing spatial proximity. Indeed the ability of DCA to distinguish correlations, that arise as a result of direct or indirect effects of nucleotide interactions, strongly increases its prediction accuracy especially in comparison with other methods such as the Mutual Information (MI).

To evaluate the performance of these DCA-based methods on RNA contact prediction, we tested them on a well curated dataset of RNA structures that we have recently established [32]. In the analysis we did not observe any significant variation among the algorithms performance for RNAs. In particular and in contrasts to results for proteins, we did not detect significant accuracy differences between mean-field and pseudo-likelihood maximization. Quite recently machine learning-based approaches have proven to astonishingly improve the prediction of protein contact maps and to considerably boost the protein 3D structure prediction [33, 34, 35]. These methods rely on the ability of deep neural networks to identify patterns in the input data using multiple levels of abstraction and have been already used to dramatically improve fields such as the computer vision and speech recognition [36, 37].

These approaches, however, are characterized by a huge number of free parameters and require big datasets of 3D structures for their training and thus cannot be easily extended to RNA structure prediction due to the limited number of available experimentally resolved structures. Here, we thus focus not on deep but on shallow Neural Networks. In particular, we construct our approach CoCoNet as combination of the mean-filed DCA approach with a shallow Convolutional Neural Network. We will demonstrate the approaches ability to improve RNA contact prediction, while keeping the number of free parameters to train the network limited to assure the generalization of its performance.

## 2 Method

### 2.1 Coevolution models

Mutations play an essential role in shaping the evolution of all biomolecules. Their large majority have a neutral effects, some of them lead to new functions while other have detrimental effects on biomolecular fitness. In the latter case the evolutionary pressure act on the biomolecules to restore their functional states favoring secondary compensatory mutations. The interactions between these mutations can be traced in the biomolecules evolution and be observed in multiple sequence alignments (MSAs) of homologous proteins or RNA. A series of co-evolutionary methods have been developed to capture these sequence variability in MSAs such as the Direct coupling analysis (DCA)[23, 24, 25] that is an inverse statistics method that are able to identified pairs of residues that co-evolved during evolutionary history and thus are likely to be in spatial adjacency in the three-dimensional structure of a protein/RNA molecule.

Let consider a sequence of nucleotide bases *σ* = *a*_1_*a*_2_*a*_3_…*a*_*L*_ of length *L* containing residues or a gap at sites 1, 2, 3, …, *L*. The probability *P* of observing this sequence in a MSA is given by the following expression is given by

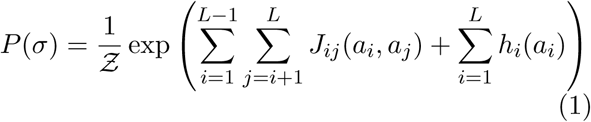

where *Ƶ* is the normalization constant (also known as partition function); *J*_*ij*_(*a*_*i*_, *a*_*j*_) are the couplings and *h*_*i*_(*a*_*i*_) are local fields. Finding a solution for equation 1 is computationally costly since the partition function scales as *𝒪*(*q*^*L*^). As a consequences, most algorithms of DCA rely on approximations. One of the most popular DCA algorithms is the mean-field (mfDCA) [25] direct-coupling analysis which shows good results for RNA[20]. It is at the same time an accurate and fast method. As numerically more complex methods such as plmDCA[38] do not lead, unlike for proteins, to improvements for RNA contact prediction [4, 20, 21] we will here focus on mfDCA.

In mfDCA the couplings are computed from the inverse of the empirical correlation matrix obtained from MSA. Let *f*_*i*_(*a*_*i*_) be single-site frequency counts of the MSA for column *i* when occupied by a nucleotide/gap *a*_*i*_, and *f*_*ij*_(*a*_*i*_, *a*_*j*_) be the pair-site frequency counts for columns *i* and *j* when occupied by nucleotide/gap *a*_*i*_ and *a*_*j*_, respectively. These quantities are computed from MSA as

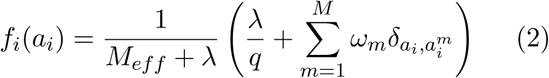

and

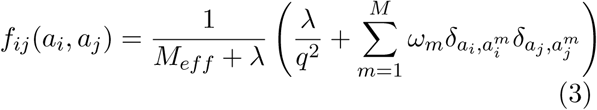

where *λ* is the pseudocount for regularizing frequency counts; *ω*_*m*_ is weight of sequence *m* which is defined as the reciprocal of the number of similar sequences for a particular sequence similarity threshold; and *M*_*eff*_ is the effective number of sequences which is the sum of sequence weights. The correlation matrix *C* has elements *C*_*ij*_ = *f*_*ij*_(*a*_*i*_, *a*_*j*_) − *f*_*i*_(*a*_*i*_)*f*_*j*_(*a*_*j*_). The couplings of the model are obtained from

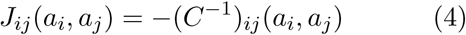

for distinct site pairs *i* and *j*. The nucleic acid pairs are scored using the direct-information that is given by

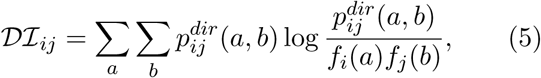

where 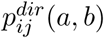 is the direct probability defined by

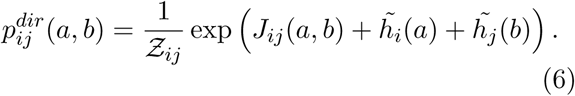

and where parameters 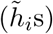 in equation 6 are obtained by requiring the direct probability marginals to be consistent with single-site frequencies of the MSA. *Ƶ*_*ij*_ is the normalization constant for 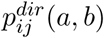. According to their DI scores, the pairs and then ranked. High-ranking pairs correspond to strongly coevolving nucleobases and thus tend to be in physical contacts in the 3D structure of the RNA molecule (true positive/ TP prediction). However, lower ranking pairs are less likely to be a real or true positive contact (TP) and more likely to be a false positive prediction (FP) not in contact in the 3D structure. It should be noted that there is no hard threshold for the DI scores, e.g. above which TP rates are high and FP rates low. Instead, there is a gradual increase of FP as one goes down the ranked pairs. Also, it should be noted that coevolution can result not only from a single native conformation but also from multiple conformations, i.e. FP can be TP in other contexts. Examples include active and inactive conformations [39] or competition of inter- and intra-contacts in homodimers [40].

### 2.2 Convolutional Neural Networks

Convolutional neural networks (CNNs) have been extensively used in the last decades in a wide range of applications that range from accurate learning of patterns in images to speech recognition [36, 37, 41]. The success of CNNs resides in their ability to identify patterns in the input data using multiple levels of abstraction through a hierarchy of different layers of convolution. These artificial networks are composed by three kinds of layers in addition to the input and output layers. The first one is the convolution layer that applies a convolution operation on the input layer, the second ones are the pooling layers that perform down-sampling operations and finally there are the fully connected layers whereby neurons are connected with all neurons in the preceding layers.

The tremendous effort devoted to the improvements of CNN architectures aims to make CNN scalable to larger and increasingly complex systems. Indeed from the simple LeNet architecture introduced in [42] consisting of three convolution, two pooling and a fully connected layers, a series of deeper CNNs that show improved performances such as AlexNet [43], ZFNet [44], GoogleNet [45] and VGGNet [46], have been introduced in the literature.

The increased level of complexity of these networks is reflected in the number of free parameters to train that range from 60k for LeNet to about 1380k for VGGNet. However, despite the accurate performances of these networks, this huge number of parameters makes the training slow and limit generalization [47].

When the training dataset is very small as for a RNA structure dataset [32], the deep network approach has to be completely ruled out to avoid overfitting and to allow reasonable generalization. For all these reasons we thus chose to employ a shallow convolutional neural network covered in the next subsection. Indeed these type of CNNs [48, 49], that have just from one to few hidden convolutional layers, while keeping good performances, are characterized by a low time training and a reduced number of free parameters.

### 2.3 Convolution on Coevolution

In order to improve contact prediction accuracy from RNA multiple sequence alignments, we here design a method called CoCoNet that is based on a combination of DCA and convolutional neural network approaches. This approach is motivated by the simple observation that contact maps of RNA are not random but instead show ordered patterns of contacts. It’s very likely that nucleotide pairs close to other pairs that are in physical contact are also true contacts themselves. CNNs are a systematic method to identify patterns from DCA contact map prediction and filter out noisy and unwanted artifacts. The architecture of our CoCoNet method is schematically depicted in Fig. 1 and is constituted by different layers.

**Figure 1:**
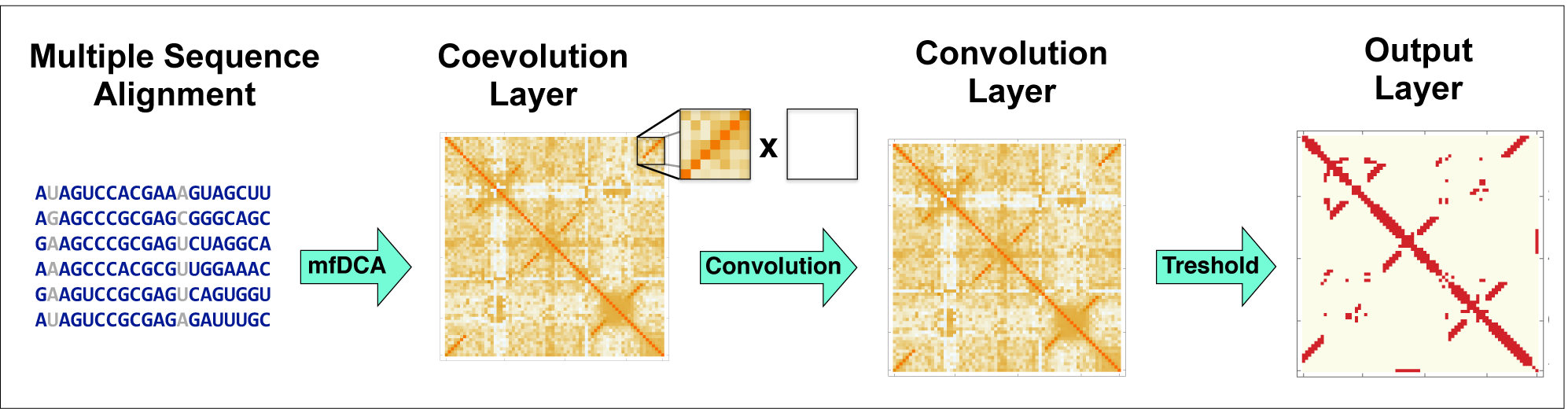
Schematic workflow of CoCoNet architecture with all different layers.

- The **input layer** is simply given by the MSA of the target RNA sequence of length *L* with its homologous.
- The first layer is the **coevolutionary layer**. In this layer the DCA scores of nucleotide-nucleotide pairs are computed using a meanfield DCA approach. This step is performed using the mean-field algorithm implementation in pydca [30]. A 2D map of size *L* × *L* is then constructed from these DCA scores assigning to each (*i, j*) pair of the target sequence the corresponding DCA score.
- The second layer is the **convolutional layer**. As a first step we perform a padding operation of size *p* = (*d* − 1)*/*2. Then a *d* × *d* filter matrix (with *d* chosen here to be equal to 3, 5 and 7) is used to perform convolution across the 2D DCA contact map obtained from the previous padded layer. This results in a new 2D contact map of size *L* in which each entry corresponds to a sort of re-weighted DCA scores.
- The **output layer** consist in selecting the *n* pairs of the previous layer map with the highest score and consider them as contact while giving a vanishing score for all the others.

### 2.4 The dataset of RNA structures

In order to train CoCoNet, we have to select a dataset of RNA structures. Here, we chose the well-curated dataset presented in [32] in which there are about seventy RNA structures of high resolution and their corresponding RFAM family of homologous RNA [50]. From all these structures we chose a subset *𝒮* of 57 entries associated to unique families in the RFAM database after discarding similar structures that belong to the same family to avoid an bais at the training state. *𝒮* is further divided in two subsets what we call *𝒮*^*H*^ and *𝒮*^*L*^ containing all entries associated to RFAM with *M*_*eff*_ greater than, and less than or equal to 70.0, since nucleotide contact prediction methods performance may depend on *M*_*eff*_ [32].

Annotations of the secondary structure has been computed using DSSR [51, 52]. The list of all PDB used in this paper and the corresponding RFAM families can be found in Table S1.

### 2.5 Learning the filter matrix

To learn the filter matrix of CoCoNet we use a gradient backpropagation algorithm. Basically, we compare the weighed contact maps for all the target sequences in our dataset that are obtained from MSAs via the coevolution plus convolutional layer with the real contact maps obtained from the PDB structures. Two nucleotides are considered as contacts in the structures if they have a pair of heavy atoms (i.e. non-hydrogen) that are less than 10 Å apart. For nucleotide pairs fulfilling this condition they are assigned a value of one in the real contact map, zero otherwise.

Given a target RNA sequence *R* belonging to the training dataset, we can define a function

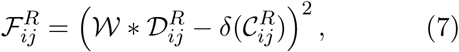

where *\** is the convolution operation between *𝒲* and *𝒟*_*ij*_ that are the filter matrix and the local *d* × *d 𝒟ℐ* scores matrix (eq. 5) centered at residue pairs (*i, j*), respectively. The delta function 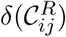 is one when nucleotide *i* and *j* are in physical contact in the PDB structure and zero otherwise.

The convolution operation can in principle be done using several filter matrices. To limit the number of free parameters, CoCoNet is designed to use a maximum of two filter matrices. Their total number range from 9, for a single 3 × 3 filter matrix up to 98 for two 7 × 7 filter matrices. When two filter matrices are used, one of them performs convolution with Watson-Crick nucleotide pairs and the other on non-Watson-Crick pairs.

The total cost function is then defined as

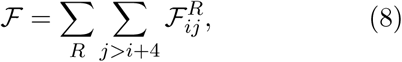

where the summation over *R* represents the summation over all the entries in the training dataset and that of *i* and *j* over all nucleotide pairs that are separated at least four nucleobases in the sequence of *R*. The cost function is minimized using Limited-memory BFGS algorithm using a standard implementation in Python’s Scipy library [53]. To ensure a strict separation of training and test data, the computation is done using a strict five-fold cross-validation with the full set randomly partitioned. The cross-validation procedure is repeated ten times and the results are obtained by averaging over all of the (ten) trials.

## 3 Results

### 3.1 Coevolutional structural features

Here, we analyze the structural patterns observed in the coevolutional layer of our network since their understanding provides insight on how CoCoNet is able to identify them and enhance nucleotide-nucleotide contact prediction. In particular, we study these structural features by investigating the average DCA scores in a 7 × 7 window around nucleotide pairs following a similar approach to the one employed in [54] for proteins. In Fig. 2.a-c we plot this average for all type of contacts according to the spatial distance *r* between the closest heavy atoms (i.e. non-hydrogen) of a nucleotide pairs. At short distance (*r* ≤ 4 Å, Fig 2.a) we clearly observe a signal corresponding to a stem structure. For this pattern the co-evolutionary scores are very strong reflecting the strong selection pressure of maintaining the corresponding secondary structure. At intermediate distance (4 < *r* ≤ 10 Å, Fig 2.b) the observed patterns are weaker and essentially are dominated by stems pairs that are in the surrounding of the target contact. Finally at distance larger than 10 Å there is essentially no signals as we can see in Fig 2.c.

**Figure 2:**
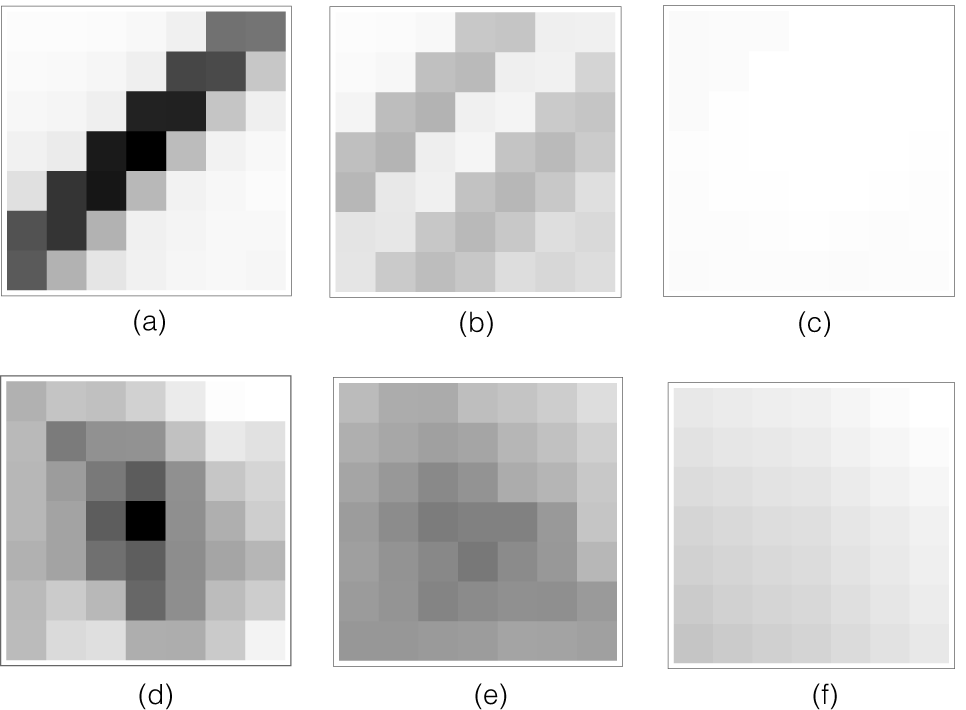
Structural features observed in the 2D coevolutionary map. Average DCA scores in a 7 × 7 window around all nucleobase pairs separated by a distance *r* ≤ 4 Å (a), 4 < *r* ≤ 10 Å (b) and *r* > 10 A (c). Here the intensity is proportional to the averaged DCA score of the corresponding element using the same color scale for (a), (b) and (c). In (d), (e) and (f) we displayed the average DCA scores in a 7 × 7 window around all 3D nucleotide pairs separated according to the same criteria *r* ≤ 4 Å, 4 < *r* ≤ 10 Å and *r* > 10 Å, respectively. Here the intensity color-scale is rescaled of a factor of about 15 when compared with (a), (b) and (c) in order to better see the patterns.

A similar pattern analysis shows when considering only nucleotide pairs that are far away from any secondary contacts, *i*.*e*. are outside a 9 × 9 window centered at any 2D contact. These patterns are shown in figures 2.d, 2.e and 2.f for distance *r* ≤ 4.0 Å, 4.0 < *r* < 10.0 Å and *r* > 10.0 Å, respectively. The first thing that we note from them is that coevolutionary signals from 3D contacts are much weaker than 2D ones: they are suppressed by a factor of about ∼ 10-20 and thus their intensity has been re-scaled accordingly to make them visible in figures 2.e-f. The patterns that we observe at short distances (2.d) has a relatively stronger signals at the middle of the windows where the 3D contact is located and tends to decrease as we move away from the center. A somewhat similar signal with a center region characterized by a stronger coevolution can be observed also at intermediate distance (2.e) even if the intensity is weaker and the pattern can be confused with the background without a further intensity rescaling (data not shown). Finally at large distance (2.f) no coevolutionary signals can be identified as expected.

### 3.2 Contact prediction accuracy

Next, we test the accuracy of our contact prediction method as a function of some neural network characteristics such as the size of the filter matrices and its architecture. We use the CoCoNet prediction scores to rank nucleotide pairs since pairs showing high scores are likely to be spatially adjacent in the three dimensional structure of an RNA molecule. To asses CoCoNet performance, we compute its positive predictive value (PPV). Figures 3 and 4 show the average PPVs as a function of the rank for all pairs (*i, j*) such that |*i* − *j*| > 4 (see Figure S1 in the supplementary material for individual RNA’s PPV) and that of tertiary contacts, respectively. Nucleotide pairs are considered as tertiary contacts if they are not secondary structure pairs and are not in a 5 × 5 windows around 2D contacts. In both cases, CoCoNet shows a significant increment of PPVs over mfDCA for almost all ranks thus indicating the ability of the convolutional layer to improve contact prediction accuracy. Although no significant difference can be observed at higher ranks (for top ∼ 5/10 nucleotide pairs) between mfDCA and CoCoNet, the predictive capacity of CoCoNet is superior than mfDCA for all ranks below that. Among the different filter sizes, the 3× 3 filter matrix performs slightly better than other filter matrices up to ranks of about hundred and slightly less beyond that limit.

**Figure 3:**
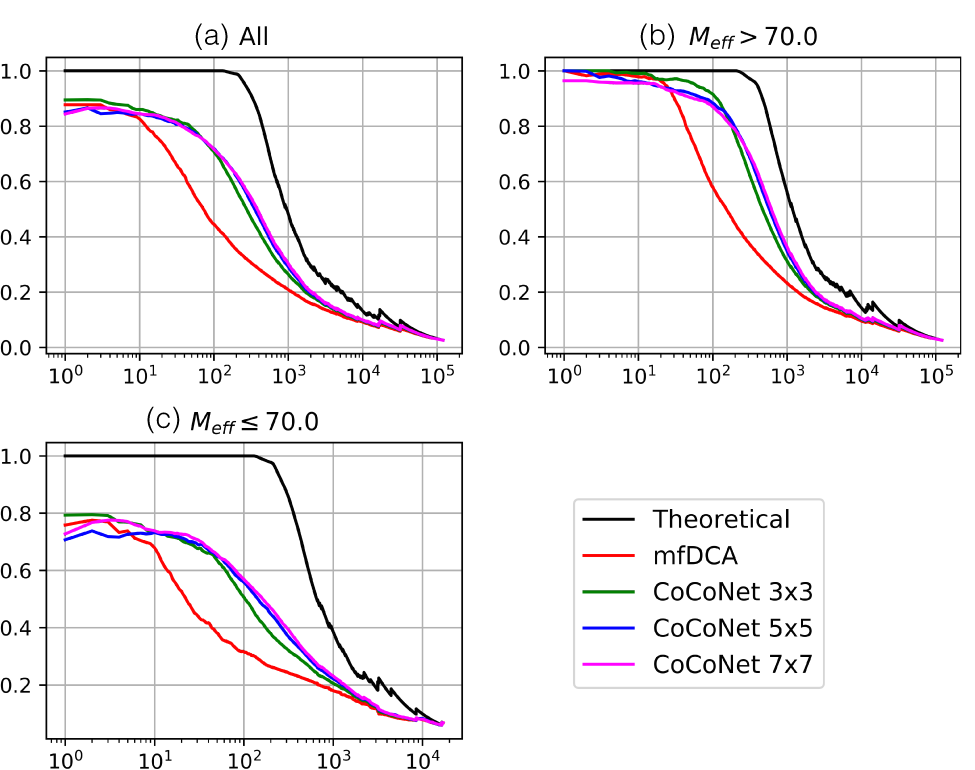
Average positive predicted value for all families in the *𝒮* (a), *𝒮*^*H*^ (b) and *𝒮*^*L*^ (c) datasets.

**Figure 4:**
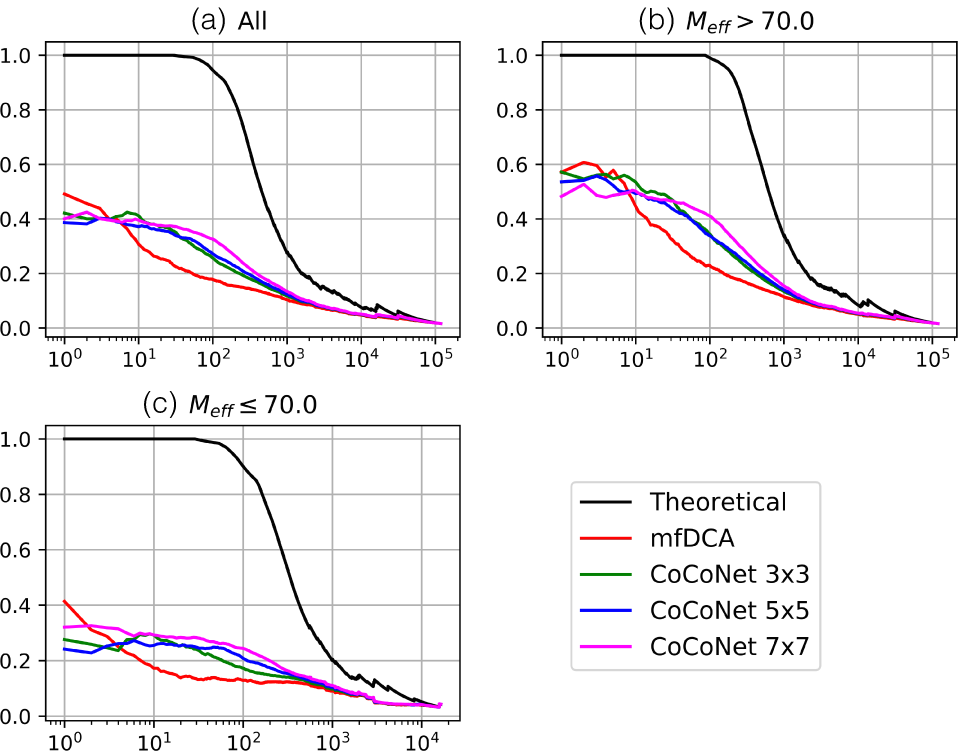
Average positive predicted value of ter-tiary contacts for all families in the *𝒮* (a), *𝒮*^*H*^ (b) and *𝒮*^*L*^ (c) datasets.

The performance of our method depend, as expected, on the effective number of homologous RNA sequences in the corresponding RFAM family of the target RNA. For families with *M*_*eff*_ > 70 the average PPVs are significantly better than those of families that have lower effective number of sequences (*M*_*eff*_ ≤ 70). This trend is consistent for both classes of contacts, i.e., all and tertiary contacts as we can see in figures 3 and 4, respectively. Nevertheless, CoCoNet outperforms mfDCA in both scenarios.

We also report the CoCoNet numerical results in table 1 where the average PPVs for top *L* contacts are displayed for different network characteristics. When all contacts are considered the performances of mean-field DCA that shows an average PPV of 45% are drastically increased to 74.5%and 77% for single and double filter versions of CoCoNet, respectively. No filter-size dependence is observed here but a slight improvement occurs by using double filter convolution with respect to the single filter ones.

**Table 1:** Average positive predicted value (⟨*PPV* ⟩) for all RNAs in the *𝒮* dataset. The first two columns indicates the number and size of filter matrices used, respectively. The third column correspond to the number of free parameters to learn. The fourth and last columns show ⟨*PPV* ⟩ at rank *L* for all and tertiary contacts, respectively. The bottom row shows the ⟨*PPV* ⟩ mean-field DCA.

| CoCoNet |  |  |  |  |
| --- | --- | --- | --- | --- |
| Filter | Filter Size | Free Param. | $\langle PPV \rangle_{ALL}$<br>(top $L$) | $\langle PPV \rangle_{3D}$<br>(top $L$) |
| 1 | 3x3 | 9 | 74.6 | 27.1 |
| 1 | 5x5 | 25 | 74.6 | 29.2 |
| 1 | 7x7 | 49 | 74.4 | 33.6 |
| 2 | 3x3 | 18 | 76.5 | 26.6 |
| 2 | 5x5 | 50 | 77.7 | 27.1 |
| 2 | 7x7 | 98 | 77.3 | 35.0 |
| Mean field DCA |  |  | 45.0 | 17.7 |

Tertiary contact prediction capability is also significantly improved by our method (see Table 1) despite the fact that their coevolutionary signals are weaker than 2D contacts as observed in section 3.1. We note here a dependence on filter matrix size since its increment is reflected by a mild increases of the PPVs 1. Still, all approaches of CoCoNet outperform vanilla mfDCA by a large margin, *e*.*g*. 35.0% vs 17.7% when using double 7 × 7 filter matrix convolution.

Finally, we also list in Table 2 the average PPVs at rank *L* for the two subsets *𝒮*^*L*^ and *𝒮*^*H*^ observing a strong improvement of the CoCoNet performances in both sets: considering all contacts in *𝒮*^*H*^CoCoNet reaches an average PPV of about 90% in comparison with 57.1% obtained from mean-field DCA. For the dataset *𝒮*^*L*^, CoCoNet’s results are even surprisingly higher reaching PPVs between 60% and 67% in comparison with 33% obtained from mean-field DCA. Similar trends are observed for tertiary contacts that are predicted with less accuracy even if their prediction remains significantly improved in both sets (see Table 2).

**Table 2:**
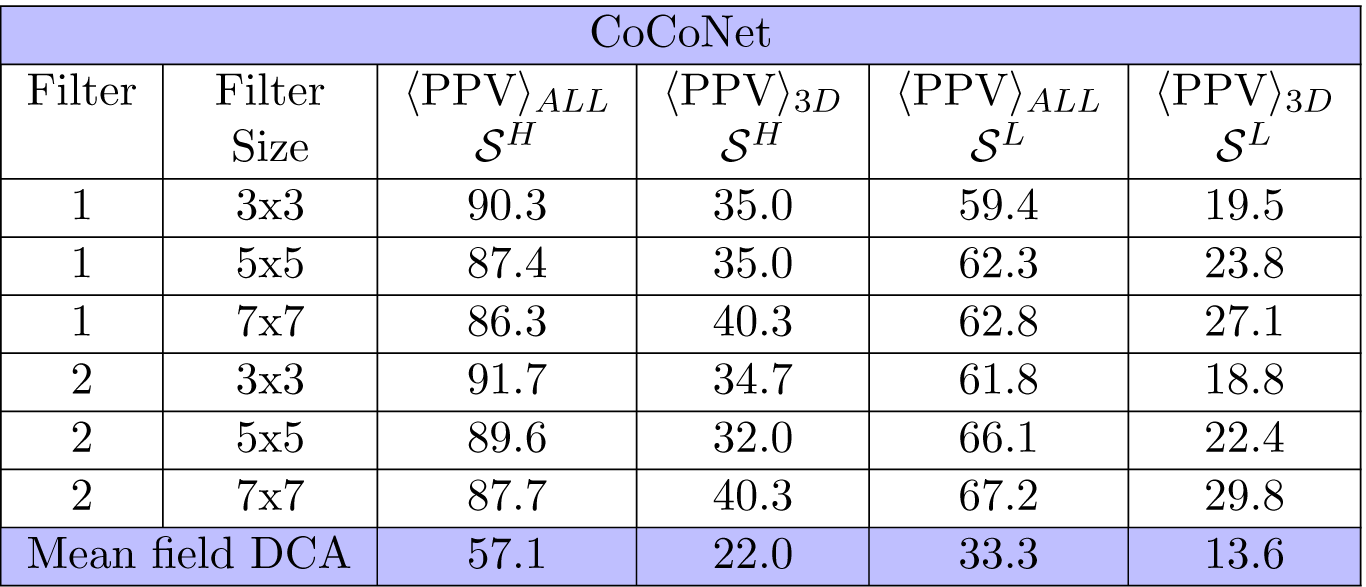
Average positive predicted values ⟨*PPV* ⟩ for for all RNAs in the *S* dataset. The first two columns indicates the number and size of filter matrices used, respectively. The third and fourth columns show ⟨*PPV* ⟩ at rank *L* in the *𝒮*^*H*^ dataset for all and tertiary contacts, respectively. Finally, the fifth and sixth columns show ⟨*PPV* ⟩ at rank *L* in the *𝒮*^*L*^ set for all and tertiary contacts,respectively. The bottom row shows averaged PPVs for mean-field DCA.

### 3.3 An example of CoCoNet application

To provide an example of the CoCoNet application we consider the aptamer domain of the Adenine Riboswitch from *Vibrio vulnificus* that has a known experimentally resolved 3D structure (see figure 5, PDB code 4TZX) [55]. This riboswitch is located in the 5’ untranslated region of the *add* adenosine deaminase mRNA and plays an important role in the translational machinery. If the adenine concentrations is high enough, the aptamer domain can bind to the adenine, induce an allosteric conformational change in the binding domains and initiate the translation. The structure consist of a three-way junction connecting three helices P1, P2, and P3 (see fig (5) with long-range three dimensional contacts occurs between P2 and P3 to stabilize the 3D structure.

**Figure 5:**
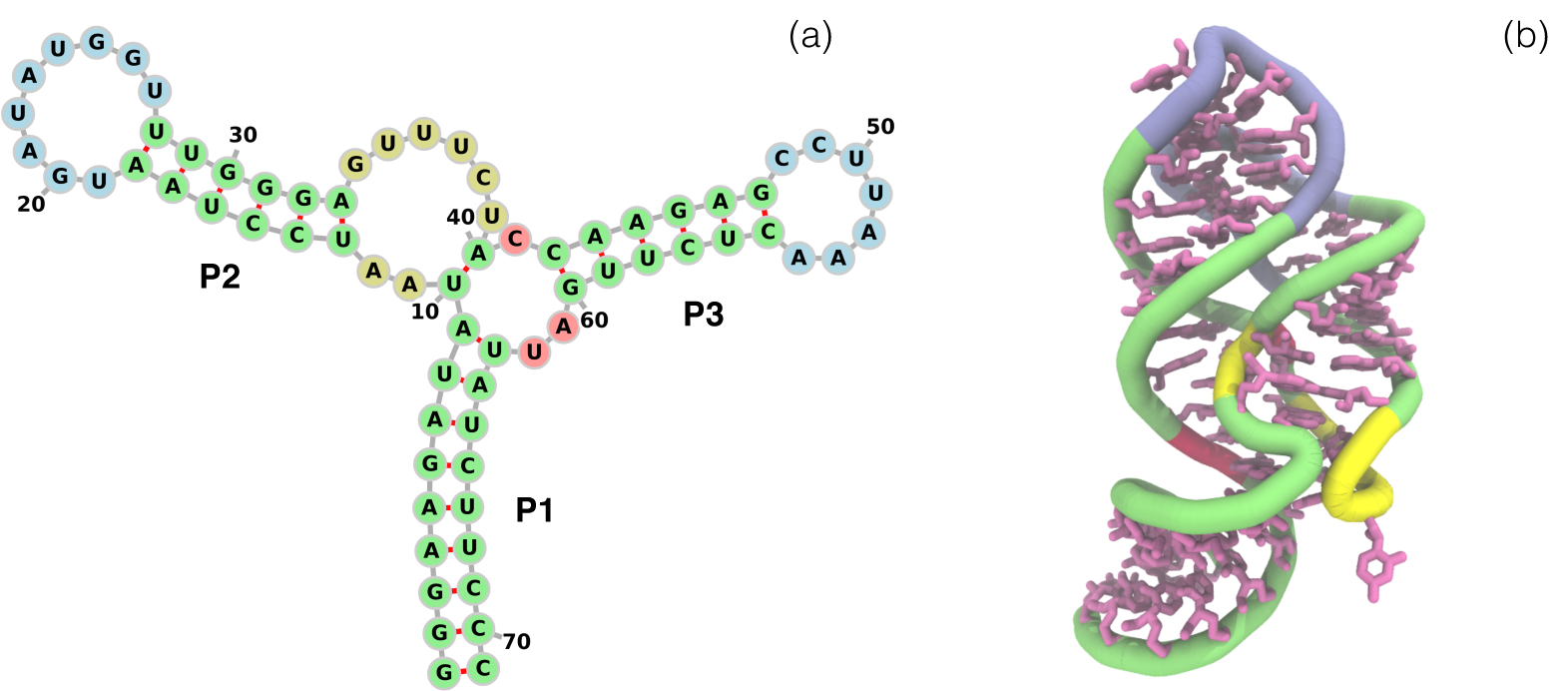
(a) Secondary and (b) tertiary structure of the *Vibrio Vulnificus* Adenine Riboswitch.

The experimental contact map of this Riboswitch is displayed in Figure 6.a where we highlight the nucleotide pairs having at least a pair of heavy atoms less than 10 Å apart. Among all these 382 contacts, the secondary structure pairs are colored in blue whereas the remaining contacts are colored in grey. Fig. 6.b display the contact map constructed by taking the top 382 mean-field DCA predicted nucleotide pairs: 38% of them are true positives (colored in green) and the rest are false positives (colored in black). Finally, fig. 6.c and 6.d represented CoCoNet predicted top 382 nucleotide pairs using 3 × 3 and 7 × 7 single filter convolution, respectively. As we can clearly see from this picture, CoCoNet (with a PVV of 60% and 67% for 3 × 3 and 7 × 7 filter size, respectively) improves the performances of mfDCA (PPV equal to 38%) substantially.

**Figure 6:**
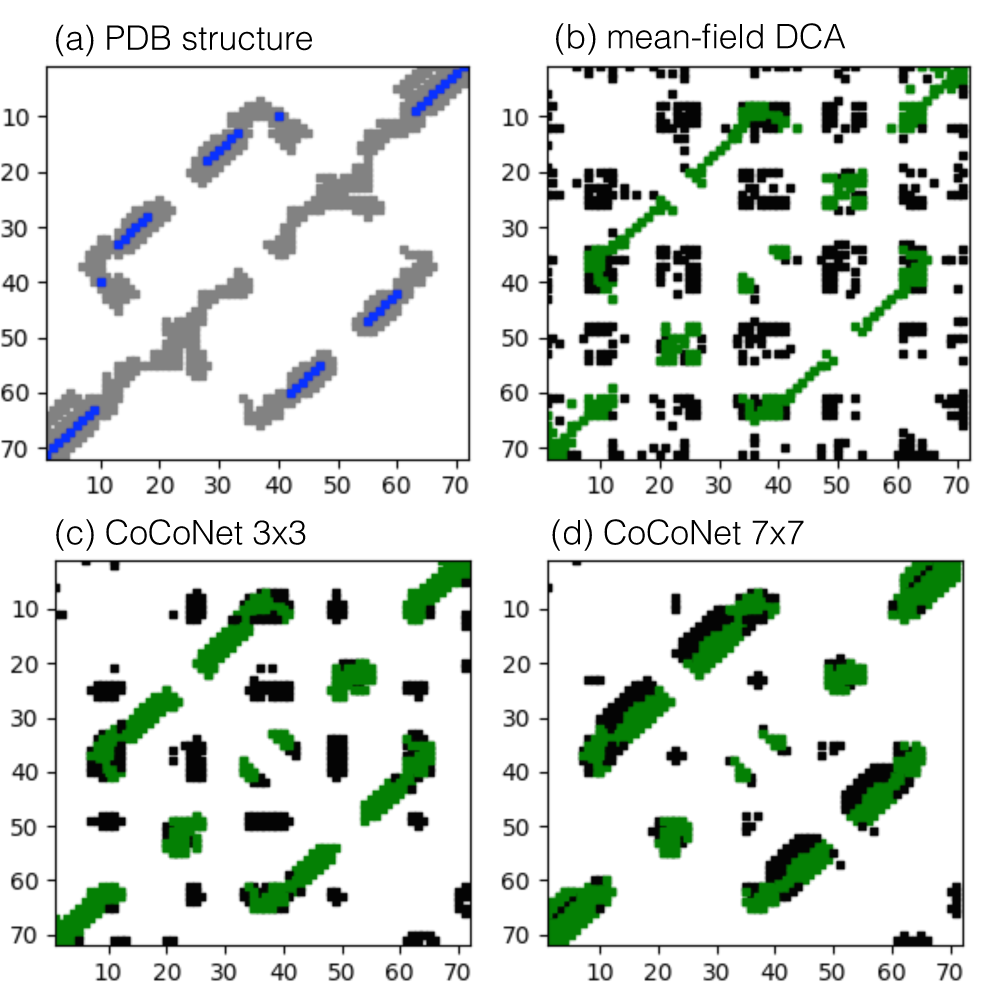
Predicted and experimental contact maps for Adenine Riboswitch from *Vibrio vulnificus* (PDB 4TZX, RFAM RF00167). (a) Contacts in the experimentally resolved PDB structure using heavy atom pair cut-off distance of 10 Å with secondary structure pairs in blue color. (b) Meanfield DCA predicted contact map with true/false positives highlighted in green/black. (c) and (d) CoCoNet predicted contact map using single 3 × 3 and 7 × 7 filter matrix respectively with the green/black color indicating true/false positives.

These contact maps clearly show the ability of CoCoNet to significantly enhance contact prediction from coevolutionary signals initially identified by DCA. The mfDCA contact map has indeed false positives scattered all over the contact map. When convolution is performed on top of co-evolution false positives are suppressed while true positives are enhanced and tend to cluster around strongly coevolving pairs. Finally, from fig. 6.c and 6.d we can also see that the clustering power of CoCoNet is enhance for large filter matrix size as already observed previously when the number of contacts considered is large enough.

## 4 Summary and conclusion

The accurate prediction of nucleotide-nucleotide contacts in RNA molecules remains an intriguing and challenging issue whose resolution could boost RNA structure prediction and to shed light on RNA fundamental properties and on its functions within the cell. Unfortunately, the limited number of resolved RNA structure prevent to use complex machine learning models couple or not with coevolutionary-based methods that recently have been successfully applied to proteins [34, 35].

In this paper we made a significant improvement in RNA contact prediction circumventing this limitation by using a combination of direct coupling analysis and a very simple convolutional neural network. Although the model has very few parameters, it is able to enhance contact prediction accuracy using limited RNA sequence data. Indeed the CoCoNet averaged PPV for a set of 57 RNAs that belong to distinct families of homologous RNA, improves the results of mean-field DCA with a PPV of 45.0% up to about 77.0% when top *L* ranked nucleotide pairs are considered. Remarkably, we observe that tertiary contact prediction is significantly improved from a PPV value of about 17.0% for the mean field DCA up to about 33.0%.

This improvement is achieved by performing convolution operation on top of coevolution and thus learning patterns of coevolving nucleotide pairs using simple filter matrices. The enhancement effect can be observed for either strong co-evolutionary signals but also for weaker ones that in principle are more easily confused with the background noise, as in the case of the 3D contacts or in the case of the homologous families with a limited number of RNA sequences.

We can explore multiple directions to further improve our method to better understand the structural properties of RNA molecules. First of all, when more 3D RNA structures will be experimentally available we could exploit more complex neural networks architecture to improve the accuracy of our method. In addition, although CoCoNet is able to enhance RNA tertiary contact prediction, their prediction accuracy remains limited and thus needs to be further improved. This is a challenging issue since as we have seen in previous sections the co-evolutionary signals are dominated by the secondary structures. Finally, it could be interesting to integrate the CoCoNet constraints in molecular modeling tools to analyze how much our improved predictions can results in more accurate structural RNA models.

## Acknowledgments

The authors gratefully acknowledge the Gauss Centre for Supercomputing e.V. (www.gauss-centre.eu) for funding this project by providing computing time through the John von Neumann Institute for Computing (NIC) on the GCS Supercomputer JUWELS at Jülich Supercomputing Centre (JSC).

